# Hexokinase 1 forms rings that constrict mitochondria during energy stress

**DOI:** 10.1101/2023.03.20.533440

**Authors:** Johannes Pilic, Benjamin Gottschalk, Benjamin Bourgeois, Hansjörg Habisch, Zhanat Koshenov, Furkan E. Oflaz, Yusuf C. Erdogan, Varda Shoshan-Barmatz, Tobias Madl, Wolfgang F. Graier, Roland Malli

## Abstract

Metabolic enzymes can adapt during energy stress, but the precise mechanisms and consequences of these adaptations remain understudied. Here, we discovered that hexokinase 1 (HK1), a key glycolytic enzyme, clusters into ring-like structures around mitochondria during energy stress. These HK1-rings constrict mitochondria at contact sites with the endoplasmic reticulum (ER) and prevent mitochondrial fission by displacing the dynamin-related protein 1 (Drp1) from mitochondrial constriction sites. Mechanistically, we identified that the lack of ATP and glucose-6-phosphate (G6P) promotes the clustering of HK1. Moreover, we found several mutations that are critical for the formation of HK1-rings. Utilizing these mutations, we could show that HK1-rings keep mitochondria connected and rewire cellular metabolism during energy stress. Our findings highlight that HK1 is a robust energy stress sensor that regulates the shape, connectivity and metabolic activity of mitochondria. Thus, the formation of HK1-rings may affect mitochondrial function in energy stress-related pathologies.

## Introduction

Enzymes are known for their catalytic activity but not for other functions. These non-catalytic functions are common. A study found that one-third of yeast enzymes had knockout-phenotypes that were restored by catalytically inactive versions of the enzymes.^1^ A widespread non-catalytic function is the reorganization of enzymes into reversible clusters upon nutrient starvation.^2^ These clusters can form large filaments that control the flux of pathways.^3^ For example, yeast glucokinase polymerizes into inactive filaments to prevent excessive glycolysis upon glucose addition.^4^ Based on these findings, we reasoned that investigating enzymes during changes in substrate availability will likely uncover new non-catalytic enzyme functions.

Here, we focused on mammalian hexokinase 1 (HK1), which catalyzes the ATP-dependent phosphorylation of glucose, because the purpose of its mitochondrial localization is not entirely clear. We used live-cell imaging to see if HK1 reorganizes during changes in glucose availability. We discovered that HK1 clusters into ring-like structures around mitochondria during glucose depletion. Using genetically encoded biosensors, we found that the formation of HK1-rings is caused by glucose depletion-induced energy stress. We designed different variants and mutants of HK1 to determine which structural motifs are crucial for the formation of HK1-rings.

## Results

### Hexokinase 1 clusters into ring-like structures during energy stress

To test how glucose availability affects the organization of HK1, we imaged HeLa cells expressing HK1-GFP and mitoDsRed, a fluorescent protein targeted to the mitochondrial matrix. HK1 was distributed homogenously around mitochondria when glucose was present (Figure 1A, left images). Upon glucose depletion, HK1 formed bright clusters throughout the mitochondrial network (Figure 1A, right images and time-lapse in Video S1A). Closer observations indicated that HK1-clusters grew in size (Figure 1B) and formed ring-like structures around mitochondria (Figure 1C) during glucose depletion. Three-dimensional reconstruction of z-stack images revealed the shape of HK1-clusters (Figure 1D and Video S1B), and that HK1 formed a closed ring around mitochondria (Figure 1E and Video S1C). HK1-rings and HK1-clusters rapidly disassembled within 1.6 ± 0.3 min (mean ± SD) upon glucose readdition (n = 10; time-lapse in Video S1D).

**Figure 1.**
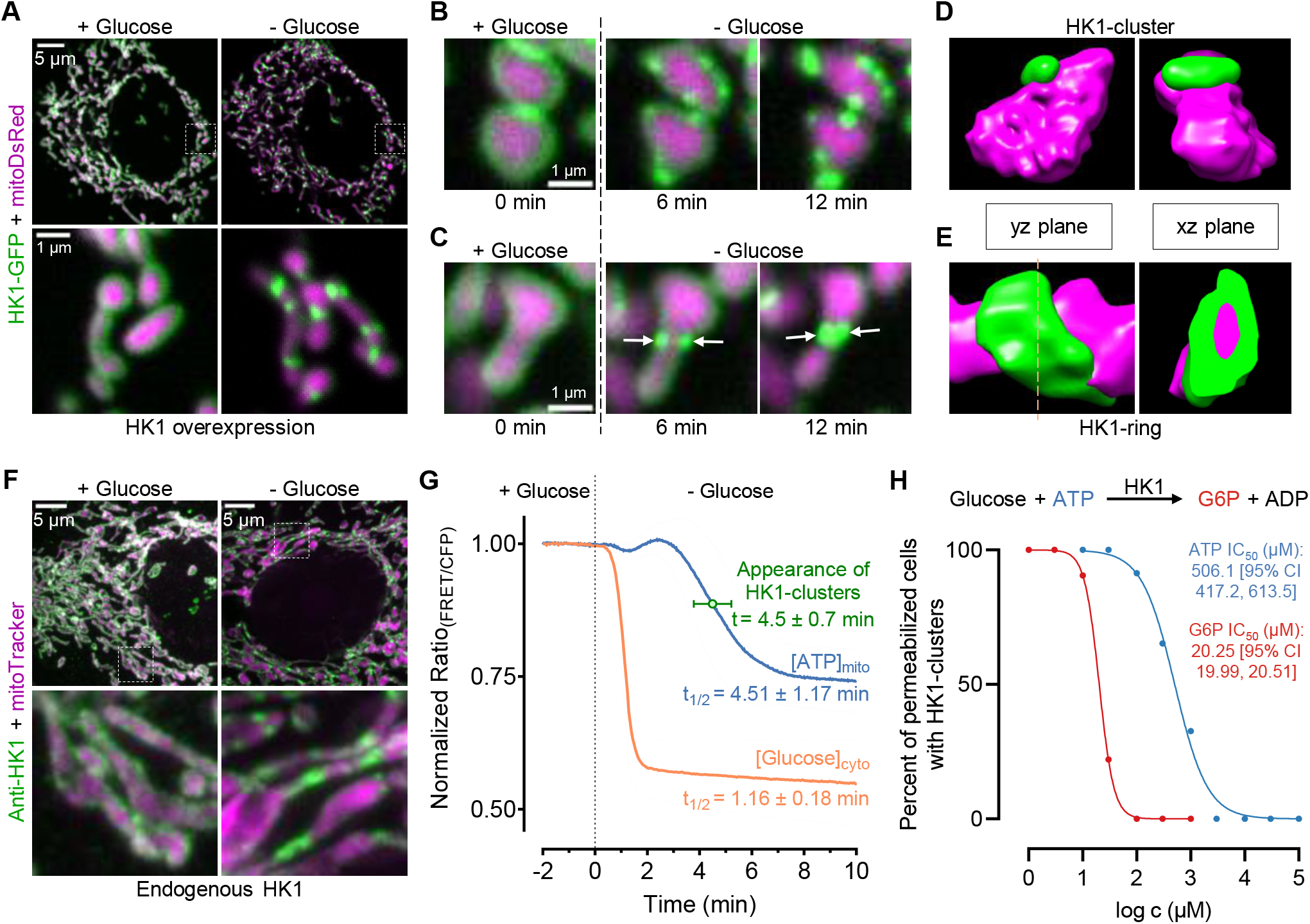
Hexokinase 1 clusters into ring-like structures during energy stress. (A) HK1 forms bright clusters when glucose is depleted. Confocal images of a HeLa cell expressing HK1-GFP and mitoDsRed with 10 mM glucose (left) and after 15 min of glucose depletion (right). The upper panels show an overview of the cell, and the dashed squares are magnified below. (B and C) HK1-clusters grow in size and form ring-like structures around mitochondria. Time-lapse images of HeLa cells expressing HK1-GFP and mitoDsRed were acquired as glucose was depleted. The maturation of multiple HK1-clusters (B) or the formation of an HK1-ring (white arrows in C) is shown. (D and E) 3D reconstructions reveal the shape of HK1-clusters and HK1-rings. Three-dimensional reconstruction of single mitochondria in HeLa cells expressing HK1-GFP and mitoDsRed. Shown are the frontal view (left panels) and side view (right panels) of an HK1-cluster (D) and an HK1-ring (E). (F) Endogenous HK1 forms ring-like structures during glucose depletion. Immunofluorescence images of MCF-7 cells stained with anti-HK1 (green) and mitoTracker (magenta) with 10 mM glucose (left) or after 30 min of glucose depletion (right). The upper panels show an overview of the cells, and the dashed squares are magnified below. (G) The appearance of HK1-clusters correlates more tightly with mitochondrial ATP than cytosolic glucose levels. Shown are curves of mitochondrial ATP levels (blue, n = 123 HeLa cells expressing mtAT1.03) and cytosolic glucose levels (orange, n = 67 HeLa cells expressing FLII12Pglu-700uDelta6) as glucose is depleted. Time-lapse images of HeLa cells expressing HK1-GFP were acquired at intervals of 5 s and used to assess the appearance of HK1-clusters (green, n = 10) as glucose is depleted. Data are presented as mean ± SD. (H) ATP and glucose-6-phosphate (G6P) disassemble HK1-clusters in a concentration-dependent manner. The mean inhibitory effects of ATP (blue, n = 46 cells) and G6P (red, n = 95 cells) on the proportion of digitonin-permeabilized HeLa cells with HK1-clusters are shown as logarithmic concentration-response curves.

Since GFP fusion constructs are prone to overexpression artifacts,^5^ we validated our findings with immunofluorescence. In line with the live-cell imaging experiments using cells expressing HK1-GFP, glucose depletion also caused the formation of endogenous HK1-clusters in MCF-7 cells (Figure 1F). To test the generality of our observations, we assessed HK1-clustering in different cell types. HK1-clusters were present in over 90% of HK1-GFP-positive HeLa cells after 20 min of perfusion with a glucose-free buffer (Figure S1A). The proportion of cells with HK1-clusters was similar in other cell types, including neuroblastoma cells (SH-SY5Y) and primary mouse embryonic fibroblasts (MEFs) (Figure S1A).

However, in rat insulinoma cells (INS-1), only 7% of HK1-GFP-positive cells formed HK1-clusters after 20 min of glucose depletion (Figure S1A). Since INS-1 cells can maintain ATP levels over long periods of glucose depletion,^6^ we hypothesized that HK1-clustering is caused by a lack of ATP and not by a lack of glucose. Therefore, we tested whether the appearance of HK1-clusters correlates more tightly with mitochondrial ATP than cytosolic glucose levels using genetically encoded fluorescent biosensors.^7,8^ In HeLa cells, HK1-clusters appeared within 4.5 ± 0.7 min after glucose depletion, which correlated well with the drop in mitochondrial ATP with a half-time of 4.51 ± 1.17 min (Figure 1G). In contrast, cytosolic glucose levels declined rapidly after glucose depletion with a half-time of 1.16 ± 0.18 min (Figure 1G). These data point towards a link between low mitochondrial ATP levels and the formation of HK1-clusters. To further test this hypothesis, we controlled intracellular ATP and glucose levels and monitored HK1-clustering in digitonin-permeabilized HeLa cells. Permeabilization of cells with a glucose-free buffer resulted in the formation of HK1-clusters (Figure S1B). In line with our assumption, the addition of ATP disassembled HK1-clusters in a concentration-dependent manner (Figure 1H, IC_50_ of 506.1 µM with 95% CI from 417.2 to 613.5 µM; images in Figure S1B). HK1-clusters remained stable in the presence of 10 mM glucose, indicating that glucose alone is insufficient to disassemble HK1-clusters (Figure S1C). Glucose-6-phosphate (G6P), the product of the hexokinase reaction, was more potent than ATP in reversing HK1-clustering (Figure 1H, IC_50_ of 20.25 µM with 95% CI from 19.99 to 20.51 µM; images in Figure S1D). The binding of G6P to the catalytically active site of HK1 is weakened by phosphate.^9,10^ Therefore, we tested whether phosphate affects HK1-clustering when G6P is present. Addition of phosphate promoted the formation of HK1-clusters, despite the presence of G6P (Figure S1E). We next tested whether ATP-binding to HK1 is important for HK1-clustering in intact cells. We designed a GFP fusion construct of a known HK1 ATP-binding mutant (G862A) with an 11-fold lower ATP-binding affinity.^11^ To induce energy stress in intact HeLa cells, glucose concentrations were gradually reduced. Cells expressing the ATP-binding mutant were approximately four times more likely than cells expressing wild-type HK1 to form clusters during perfusion with 0.33 mM glucose (Figure S1F). These data suggest that the lack of ATP promotes HK1-clustering.

### HK1-rings constrict mitochondria at ER-contact sites

We observed that mitochondria are constricted at positions of HK1-rings (Figure 2A, left panel). To assess the degree of mitochondrial constriction, we coexpressed HK1-GFP with mitoDsRed and measured the diameter of the mitochondrial matrix at positions of HK1-rings. The diameter of the mitochondrial matrix was 0.27 ± 0.05 µm (mean ± SD) with glucose, significantly decreased to a resolution-limited minimum of 0.21 ± 0.02 µm at HK1-rings without glucose, and returned to basal levels of 0.26 ± 0.03 µm after HK1-rings disassembled upon glucose readdition (Figure 2A, right graph). Since actin contributes to the constriction of mitochondria,^12,13^ we tested whether inhibition of actin polymerization with cytochalasin D before and during glucose depletion affects the formation of HK1-rings. To visualize the actin distribution relative to HK1, we coexpressed HK1-GFP with mCherry-Actin. Under basal conditions, we observed a dense network of actin fibers that disassembled upon perfusion with cytochalasin D (Figure S2A). Inhibition of actin polymerization did not affect the localization of HK1 in the presence of glucose or the formation of HK1-rings after glucose depletion. (Figure S2A). These findings suggest that actin is not required for the formation of HK1-rings.

**Figure 2.**
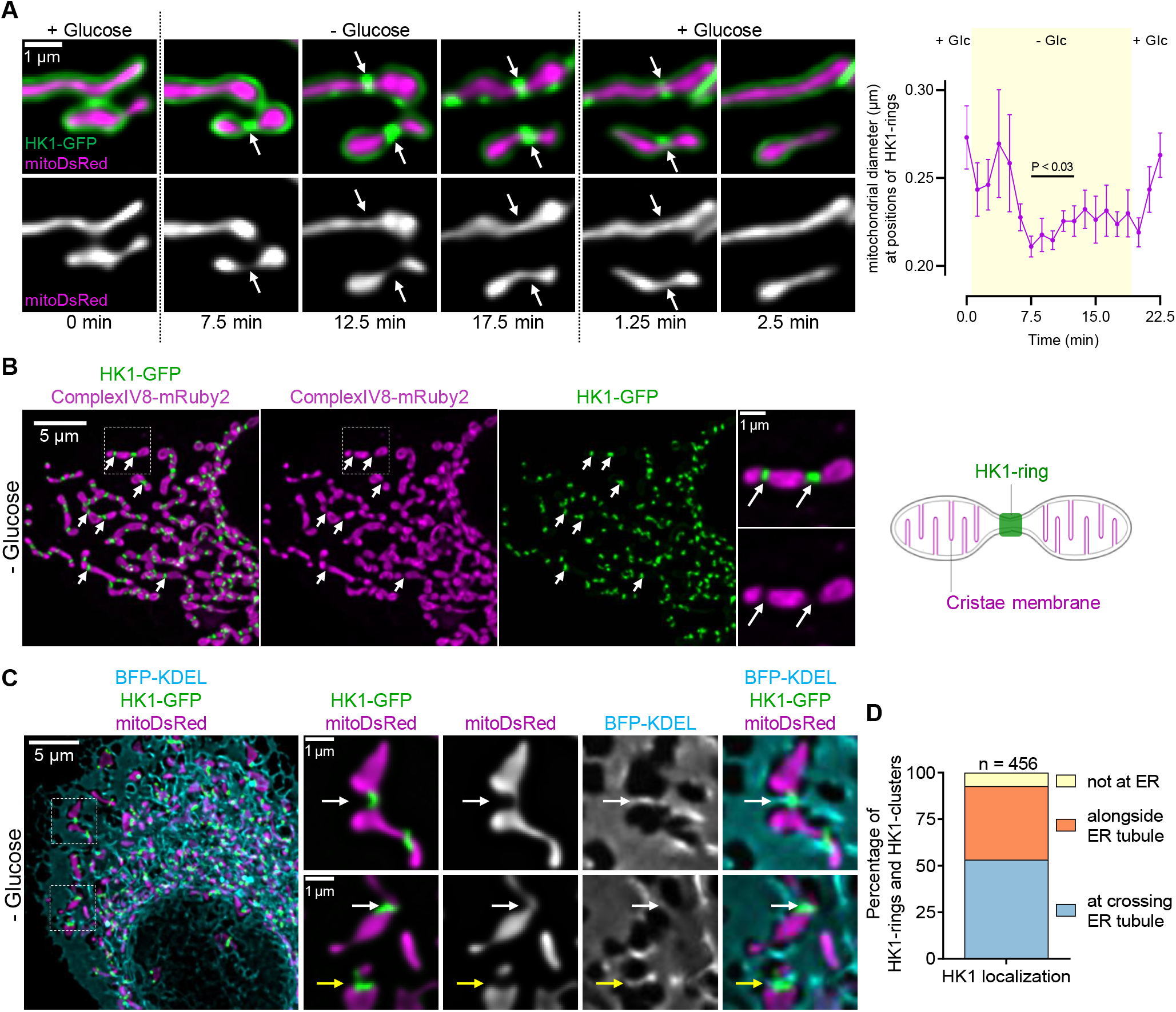
HK1-rings constrict mitochondria at ER-contact sites. (A) HK1-rings constrict mitochondria. Time-lapse images of a HeLa cell expressing HK1-GFP and mitoDsRed were acquired as glucose was removed and readded (left panel). Images are maximum intensity projections of z-stacks (6 sections, spaced 0.2 µm apart). Arrows indicate sites of mitochondrial constriction at positions of HK1-rings. The diameter of the mitochondrial matrix (mean ± SEM, n = 7) was measured at positions of HK1-rings (right graph). The difference between basal and glucose-depleted mitochondrial diameter was evaluated using one-way ANOVA with Dunnett post hoc test. (B) Cristae membranes are absent at positions of HK1-rings. Confocal images of a HeLa cell expressing HK1-GFP and ComplexIV8-mRuby2 (cristae membrane) after 15 min of glucose depletion (left panel). Images are maximum intensity projections of z-stacks (51 sections, spaced 0.2 µm apart). Arrows indicate positions of HK1-rings. Dashed squares are magnified on the right side of the overview images. Cartoon shows that cristae membranes are displaced at positions of HK1-rings (right). (C) HK1-rings are located at ER contact sites. Confocal images of a HeLa cell expressing HK1-GFP, mitoDsRed, and BFP-KDEL (ER) after 15 min of glucose depletion. The left image shows an overview of the cell, and the dashed squares are magnified on the right side. White arrows point to positions of HK1-rings, and yellow arrows to HK1-clusters. (D) The majority of HK1-rings and HK1-clusters are localized at the ER. Bar graph shows the percentage of HK1-rings and HK1-clusters that colocalize with the ER membrane after 15 min of glucose depletion (n = 456).

We assumed that the constriction of mitochondria by HK1-rings reduces the density of cristae membranes at positions of HK1-rings. To visualize such changes in mitochondrial ultrastructure in living cells, we coexpressed HK1-GFP with ComplexIV8-mRuby2, a cristae membrane marker. Cristae membranes were absent at positions of HK1-rings (Figure 2B). To confirm the presence of mitochondrial matrix in these cristae-free regions, we additionally expressed mitoBFP, a blue fluorescent mitochondrial matrix marker (Figure S2B). Colocalization analysis confirmed that HK1-rings colocalized significantly less with the cristae membrane than with the mitochondrial matrix (Figure S2C). These data indicate that cristae membranes are displaced at positions of HK1-rings.

Since the ER is known to be involved in mitochondrial constriction,^14,15^ we tested whether HK1-rings are localized at the ER using the marker BFP-KDEL. Indeed, HK1-rings colocalized with ER tubules (Figure 2C, white arrows; 3D animation in Video S2A). HK1-clusters of neighboring mitochondria were frequently observed in contact (Figure S2D). ER tubules were found between HK1-cluster contact sites (Figure 2C, yellow arrows; 3D animation in Video S2B). More than 90% of HK1-clusters and HK1-rings colocalized with crossing and adjacent ER tubules (Figure 2D). These data indicate that the ER is likely responsible for the positioning of HK1 during energy stress.

### HK1-rings displace Drp1 and prevent mitochondrial fission during energy stress

Drp1 is a critical player in mitochondrial fission and is known to constrict mitochondria.^16^ Therefore, we tested whether HK1-rings colocalized with mCherry-Drp1 and monitored their subcellular localization during glucose depletion. The number of Drp1-clusters decreased when glucose was depleted, while HK1-rings were formed (Figure 3A, images in Figure S3A). During glucose depletion, expression of HK1 significantly reduced the number of Drp1-clusters (Figure 3A, images in Figure S3A). The majority of HK1-rings and Drp1-clusters did not colocalize, and less than 20% of HK1-rings and Drp1-clusters were directly adjacent (Figure 3B, representative image in Figure 3SB). Glucose readdition increased the number of Drp1-clusters while disassembling HK1-rings (Figure 3A). We found that Drp1 clustered and triggered mitochondrial fission only at a few sites where HK1-rings disassembled (Figure 3C). HK1-rings appeared to be bulky structures. Therefore, we tested whether the formation of HK1-rings is associated with the displacement of other outer mitochondrial membrane proteins. For this purpose, we coexpressed HK1-GFP with mTagBFP2-TOMM20-N-10, a marker of the outer mitochondrial membrane, and imaged their distribution during glucose depletion. HK1-rings displaced the marker during glucose depletion (Figure 3D). Together, these findings indicate that at positions of bulky HK1-rings, Drp1 is unable to carry out fission during energy stress.

**Figure 3.**
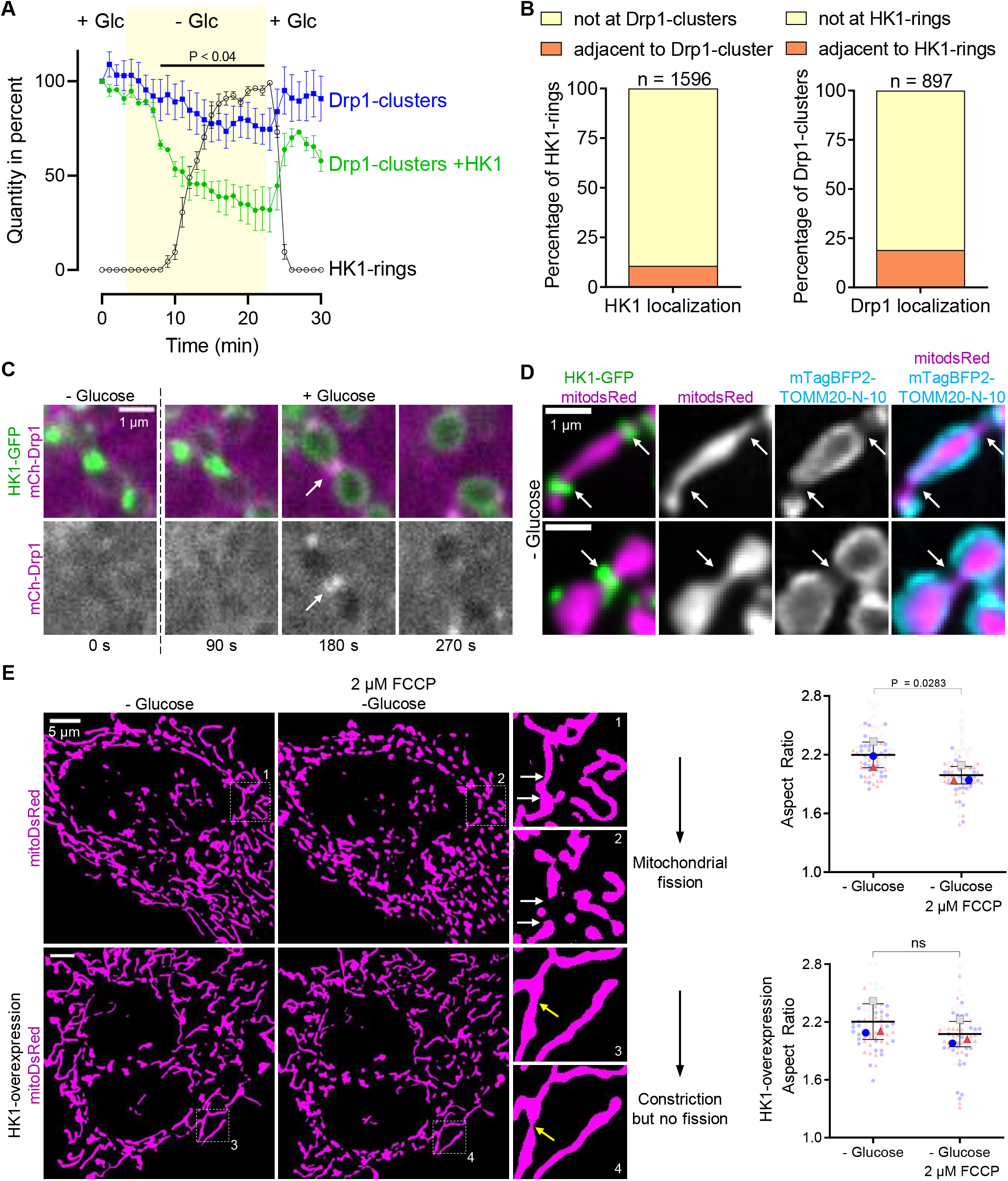
HK1-rings displace Drp1 and prevent mitochondrial fission during energy stress. (A) HK1-rings reduced the number of Drp1-clusters during energy stress. Curves show the percentage of Drp1-clusters in HeLa cells (blue) and in HeLa cells that express HK1-GFP (green) relative to the percentage of HK1-rings (black) as glucose is removed and readded. Two-tailed unpaired t-test was used for statistical analysis between Drp1-clusters and Drp1-clusters +HK1. Data are presented as mean ± SEM. (B) The majority of HK1-rings and Drp1-clusters did not colocalize. Bar graphs show the percentage of HK1-rings (n = 1596) that were adjacent to Drp1-clusters (left) and the percentage of Drp1-clusters (n = 897) that were adjacent to HK1-rings (right) after 15 min of glucose depletion. (C) HK1-ring disassembly allowed Drp1-clusters to move into constriction sites and induce mitochondrial fission. Time-lapse images of a glucose-depleted HeLa cell expressing HK1-GFP and mCherry-Drp1 were acquired at intervals of 90 s as 10 mM glucose was readded. Arrow indicates the position of an HK1-ring. (D) HK1-rings displace proteins of the outer mitochondrial membrane. Confocal images of a HeLa cell expressing HK1-GFP, mitoDsRed, and mTagBFP2-TOMM20-N-10 after 15 min of glucose depletion. Images are maximum intensity projections of z-stacks (35 sections, spaced 0.2 µm apart). Arrows indicate positions of HK1-rings. (E) HK1-rings prevent FCCP-induced mitochondrial fission during energy stress. Thresholded images of mitoDsRed signal in HeLa cells (top panel) and in HeLa cells that overexpress HK1-GFP (bottom panel). Glucose was depleted for 30 min before imaging (leftmost images). The middle images show the cells after perfusion with 2 µM FCCP for 20 min. Numbered dashed squares are magnified on the right side of the overview images. White arrows indicate mitochondrial fission, and yellow arrows indicate mitochondrial constriction. The beeswarm SuperPlot represents each cell with a color-coded dot according to the experimental day (right). Two-tailed paired t-test was used for statistical analysis (n = 3). Data are presented as mean ± SD.

HeLa cells expressing HK1-GFP appeared to be protected from mitochondrial fission (Video S3A). To challenge the protective effect of HK1-rings against mitochondrial fission, we compared the mitochondrial morphology of glucose-depleted HeLa cells with and without expression of HK1-GFP during perfusion with carbonyl cyanide-p-trifluoromethoxyphenylhydrazone (FCCP), a potent inducer of mitochondrial fission. Changes in mitochondrial morphology were determined with the ratio of the longest and shortest axes of a mitochondrial matrix marker (aspect ratio, AR). Low AR implies that mitochondria are fragmented, whereas high AR implies that mitochondria are elongated. As evidenced by the significant reduction of AR, FCCP treatment induced mitochondrial fission in HeLa cells that did not express HK1-GFP (Figure 3E, upper panel). In contrast, FCCP treatment did not induce mitochondrial fission in HK1-GFP-expressing HeLa cells (Figure 3E, lower panel). These data indicate that HK1-rings prevent FCCP-induced mitochondrial fission during energy stress.

### Structural features dictate the formation of HK1-clusters

We sought to identify which structural features are important for the clustering of HK1. First, we performed glucose depletion experiments with other mitochondrial hexokinases to test whether clustering is specific to HK1. HK2, which has 73% sequence homology and a similar structure to HK1, did not form clusters during glucose depletion (Figure S4A). The recently discovered HKDC1, which arose from a gene duplication event of HK1 and has 71% sequence homology to HK1,^17^ showed a moderate ability to form clusters during glucose depletion (Figure S4A). These data indicate that changes in the structure of the enzyme impact the ability to form clusters.

The HK1 enzyme consists of two structurally similar halves: the catalytically inactive N-terminal and the active C-terminal half (Figure 4A). In contrast, HK2 has two catalytically active halves.^18^ To identify which half is essential for HK1-clustering, we designed two chimera constructs of HK1 and HK2 fused to GFP: HK1N-HK2C and HK2N-HK1C (Figure 4B). To assess these constructs, we compared the percentage of cells that displayed cluster formation during glucose depletion. After 10 min of perfusion with a glucose-free buffer, HK1-clusters were present in 68% of cells expressing HK1-GFP. In contrast, cells expressing the HK2N-HK1C-GFP chimera did not show cluster formation under the same conditions (Figure 4B, images in Figure S4B). However, 16% of HK1N-HK2C-GFP-expressing cells showed clusters in response to glucose depletion (Figure 4B and Figure 4C). These data demonstrate that the C-terminal half of HK1 is not essential but supports cluster formation, while the N-terminal half of HK1 is essential for cluster formation upon glucose depletion.

**Figure 4.**
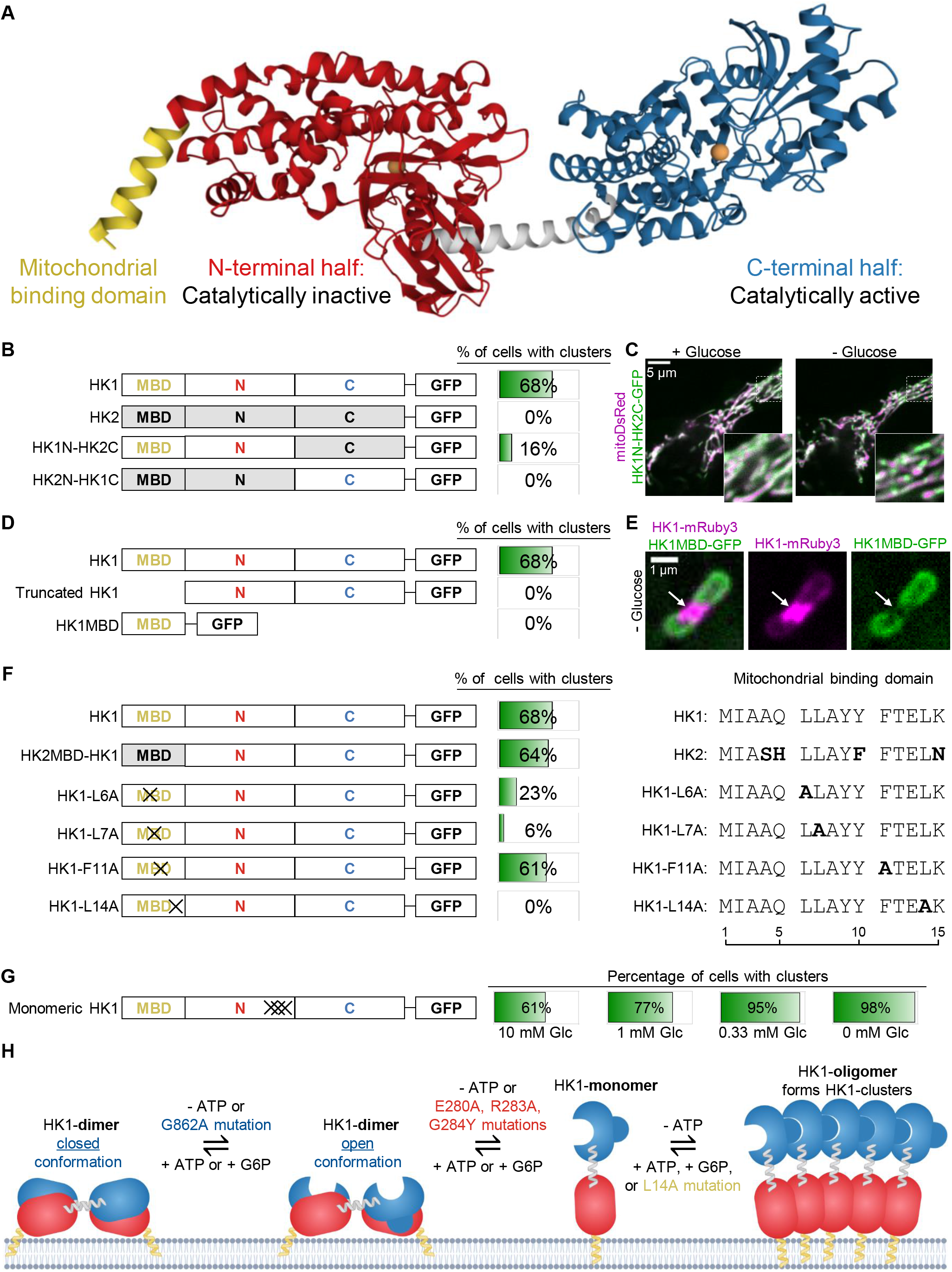
Structural features dictate the formation of HK1-clusters. (A) X-ray structures of full-length rat HK1 (PDB accession no. 1BG3). HK1 contains a mitochondrial binding domain (yellow), a catalytically inactive N-terminal half (red), and a catalytically active C-terminal half (blue). Glucose is shown as orange balls. (B) The N-terminal half of HK1 is essential for the formation of HK1-clusters. Domain organization of HK1, HK2, and chimera constructs are shown on the left next to the percentages of HeLa cells with clusters after 10 min of glucose depletion. HK1 (n = 110), HK2 (n = 57), HK1N-HK2C (n = 81), HK2N-HK1C (n = 40). MBD, mitochondrial binding domain; N, N-terminal half; C-terminal half. (C) The C-terminal half of HK1 is not essential but supports cluster formation. Confocal images of a HeLa cell expressing HK1N-HK2C-GFP and mitoDsRed with 10 mM glucose (left) and after 10 min of glucose depletion (right). Dashed squares in the overview images are magnified in the bottom right corner. (D) The MBD is necessary for the formation of HK1-clusters. Domain organization of HK1, truncated HK1, and HK1MBD are shown on the left next to the percentages of HeLa cells with clusters after 10 min of glucose depletion. HK1 (n = 110), truncated HK1 (n = 10), HK1MBD (n = 97). MBD, mitochondrial binding domain; N, N-terminal half; C, C-terminal half. (E) HK1MBD-GFP was absent at HK1-rings. Confocal images of a HeLa cell expressing HK1MBD-GFP and HK1-mRuby3 after 20 min of glucose depletion. Arrow indicates the position of an HK1-ring. (F) Leucine residues within the MBD are crucial for the formation of HK1-clusters. Domain organization of HK1, HK2MBD-HK1, and mutants in the MBD are shown on the left next to the percentages of HeLa cells with clusters after 10 min of glucose depletion. HK1 (n = 110), HK2MBD-HK1 (n = 33), HK1-L6A (n = 31), HK1-L7A (n = 33), HK1-F11A (n = 28), HK1-L14A (n = 26). Residues in the MBD of constructs that differ from HK1 are highlighted in bold (right). MBD, mitochondrial binding domain; N, N-terminal half; C, C-terminal half. Crosses indicate mutated positions. (G) Monomeric HK1 forms clusters even in the presence of glucose. Domain organization of monomeric HK1 (E280A, R283A, and G284Y) is shown on the left next to the percentage of HeLa cells with clusters of monomeric HK1 (n = 44) as glucose was gradually depleted in intervals of 7 min. MBD, mitochondrial binding domain; N, N-terminal half; C, C-terminal half. Crosses indicate mutated positions. (H) Model of HK1-clustering. Lack of ATP or mutation in the ATP-binding site favors an open conformation of the C-terminal half and is an initial step of HK1-clustering. Further depletion of ATP or mutation in the dimeric interface leads to dimer-monomer-transition. Monomeric HK1 forms new oligomers that assemble into HK1-clusters. HK1-clustering is reversed by ATP or G6P.

The N-terminal half of HK1 contains a hydrophobic mitochondrial binding domain (MBD) of 15 amino acids (Figure 4A). We tested whether truncated HK1, lacking the MBD,^19^ can form clusters during glucose depletion. The truncated HK1 was localized to the cytosol and did not form clusters during glucose depletion (Figure 4D, images in Figure S4B), indicating that the MBD or a high local concentration of HK1 is needed for the formation of HK1-clusters. To assess whether the MBD alone is sufficient for clustering, we fused the MBD of HK1 to GFP (HK1MBD-GFP) and expressed the construct in HeLa cells. HK1MBD-GFP was homogeneously distributed around mitochondria and did not form clusters during glucose depletion (Figure 4D, images in Figure S4B). To test whether the MBD can accumulate at positions of HK1-rings, we imaged HK1-mRuby3 and HK1MBD-GFP in cells during glucose depletion. HK1MBD-GFP was absent at HK1-rings (Figure 4E), indicating that the MBD alone is insufficient to integrate into HK1-rings.

Next, we sought to identify which residues of the MBD are important for HK1-clustering. Therefore, we designed a chimera construct of HK1 and the MBD of HK2: HK2MBD-HK1-GFP and monitored HK1-cluster formation during glucose depletion. The MBD of HK2 differs in residues 4, 5, 10, and 15 from the MBD of HK1 (Figure 4F). The percentage of cells with clusters was comparable in cells expressing HK2MBD-HK1-GFP and cells expressing HK1-GFP (Figure 4F, images in Figure S4C). It has been shown that replacing hydrophobic or charged residues with alanine can make membrane-curving proteins defective in their ability to tubulate membranes.^20,21^ Therefore we designed four constructs of HK1 fused to GFP with mutations in the MBD: HK1-L6A, HK1-L7A, HK1-F11A, and HK1-L14A. These mutations in the MBD did not prevent the mitochondrial localization of the constructs (Figure S4C). 61% of HK1-F11A-GFP-expressing cells, 23% of HK1-L6A-GFP-expressing cells, and only 6% of HK1-L7A-GFP-expressing cells were able to form HK1-clusters during glucose depletion (Figure 4F, images in Figure S4C). Cells expressing HK1-L14A-GFP did not show HK1-cluster formation upon glucose depletion (Figure 4F, images in Figure S4C). These data suggest that leucine residues within the MBD are crucial for the formation of HK1-clusters in response to energy stress.

Finally, we tested whether the oligomeric state of HK1 is important for HK1-clustering. Although HK1 is monomeric in solution,^22^ it is considered an oligomer of up to four subunits when bound to the outer mitochondrial membrane.^23,24^ The biological significance of monomeric HK1 is unknown. Therefore, we designed a GFP fusion construct of a known monomeric HK1 triple mutant (E280A, R283A, and G284Y).^25^ These mutations are located at the dimer interface and prevent dimerization of the enzyme.^25^ In 61% of cells expressing monomeric-HK1-GFP, clusters were found even in the presence of glucose, and glucose depletion further increased this proportion to 98% (Figure 4G, images in Figure S4D), indicating that monomeric HK1 favors the formation of HK1-clusters.

### HK1-rings keep mitochondria connected and rewire cellular metabolism

To test the functional consequences of HK1-rings formation, we focused on two established mutants, HK1-ATPmut, which favors ring formation, and HK1-L14A, which cannot form rings during energy stress. First, we challenged the protective effect of HK1-rings against mitochondrial fission, by comparing the mitochondrial morphology of glucose-depleted HeLa cells expressing HK1-ATPmut-GFP or HK1-L14A-GFP during perfusion with FCCP. FCCP treatment induced mitochondrial fragmentation in HeLa cells expressing HK1-L14A-GFP (Figure 5A, upper panel). In contrast, FCCP treatment did not induce mitochondrial fragmentation in HK1-ATPmut-GFP-expressing HeLa cells (Figure 5A, lower panel). These data indicate that single-point mutations in HK1 can alter mitochondrial dynamics during energy stress.

**Figure 5.**
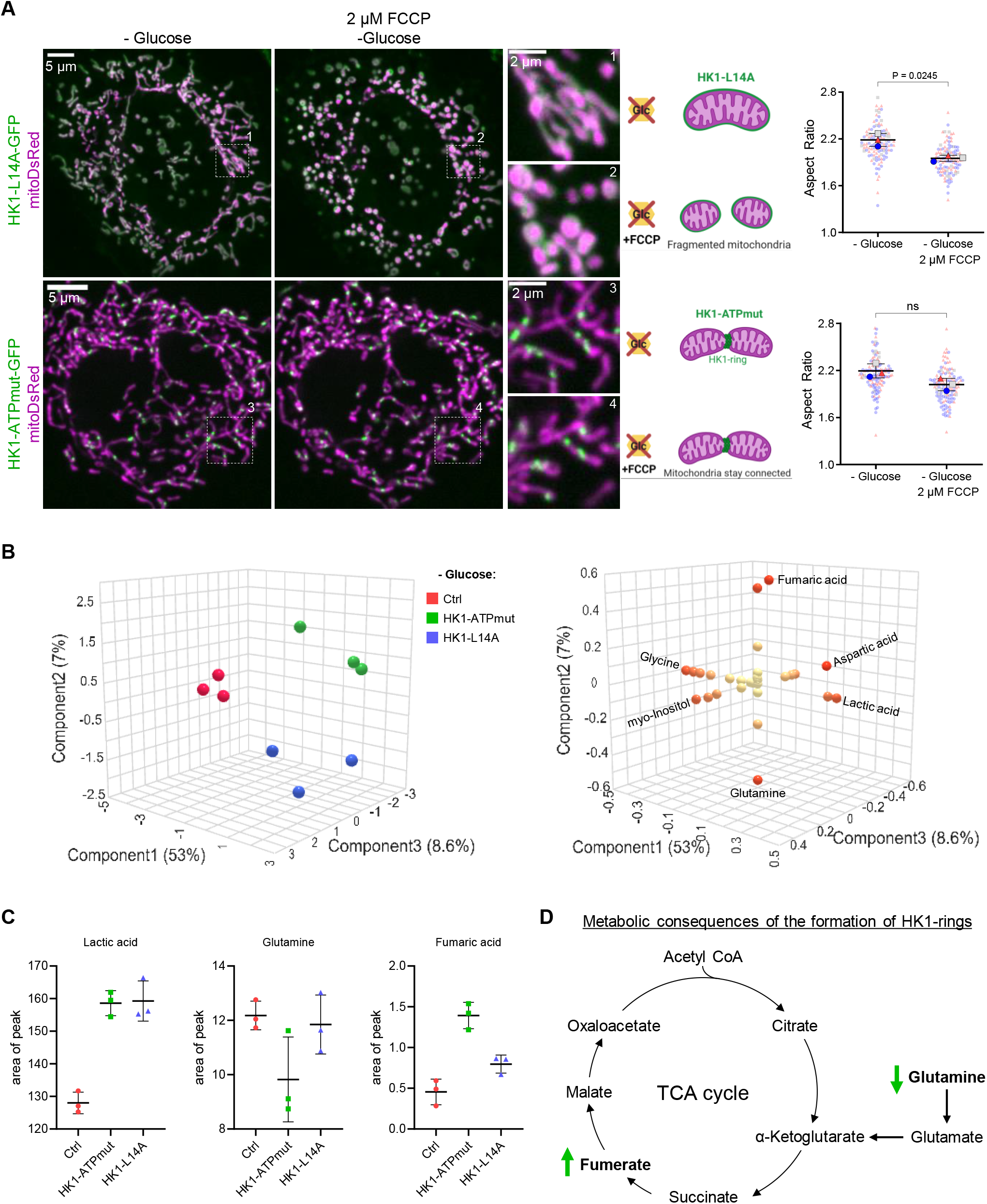
HK1-rings keep mitochondria connected and rewire cellular metabolism. (A) Single point mutations in HK1 alter mitochondrial dynamics during energy stress. Confocal images of HeLa cells expressing mitoDsRed with HK1-L14A-GFP (top panel) and HeLa cells that express mitoDsRed with HK1-ATPmut-GFP (bottom panel). Glucose was depleted for 30 min before imaging (leftmost images). The middle images show the cells after perfusion with 2 µM FCCP for 20 min. Numbered dashed squares are magnified on the right side of the overview images. The beeswarm SuperPlot represents each cell with a color-coded dot according to the experimental day (right). Two-tailed paired t-test was used for statistical analysis (n = 3). Data are presented as mean ± SD. (B) HK1-rings rewire mitochondrial metabolism. Untarget NMR metabolomics sPLS-DA 3D score plot (left): Component 1 of 53%, Component 2 of 7%, and Component 3 of 8.6%. 3D loading plot indicates most distinctive metabolites (right). (C) Graphs show changes in the area of peaks of selected metabolites. (D) Proposed model for rewiring of cellular metabolism by HK1-rings. Green arrows indicate consequences in metabolite levels by the formation of HK1-rings.

Next, we used untargeted NMR metabolomics to examine whether HK1-rings affect cellular metabolites in glucose-starved HeLa cells. Sparse partial least squares-discriminant analysis (sPLS-DA) was used to identify differences between non-transfected control cells and cells transfected with HK1-ATPmut or HK1-L14A. The sPLS-DA score plot demonstrates a clear separation of metabolic profiles among the three groups, with Component 1 accounting for 53%, Component 2 for 7%, and Component 3 for 8.6% (Figure 5B, left). sPLS-DA was used to determine the most distinctive metabolites between the groups (Figure 5B, right and Figure S5). Compared to control cells, cells transfected with HK1-ATPmut or HK1-L14A had increased lactate levels (Figure 5C), suggesting that overexpression of HK1 mutants promotes lactate production. In cells transfected with HK1-ATPmut, we measured reduced glutamine and increased fumarate levels compared to control cells and cells transfected with HK1-L14A (Figure 5C), indicating that HK1-rings favor a shift in metabolism to glutaminolysis and increased TCA cycle activity (Figure 5D).

## Discussion

Here, we discovered that HK1 forms rings that constrict mitochondria during energy stress. HK1-rings are localized at contact sites with the ER and prevent Drp1-mediated mitochondrial fission. Prevention of mitochondrial fission by HK1-rings likely preserves the integrity of mitochondria during energy stress.

Based on our findings, we present a model for the formation of HK1-clusters (Figure 4H): We propose that depletion of ATP is the first step in the formation of HK1-clusters. It is not the lack of glucose but the depletion of ATP that triggers HK1-clustering. This mechanistic insight likely explains why HK1-clustering was not discovered earlier under similar conditions. One group reported that glucose depletion did not affect the subcellular location of HK1.^26,27^ This could be due to low spatial resolution or the use of cell types that can maintain ATP levels during glucose depletion.^6^ Interestingly, glucose alone cannot stabilize a closed conformation of the catalytically active C-terminal half.^28^ A closed conformation of the C-terminal half requires ATP or G6P,^28^ which led us to believe that an open conformation of the C-terminus favors HK1-clustering. Supporting this assumption, we demonstrated that a point mutation in the ATP-binding site promotes the formation of HK1-clusters. Our data also indicate that G6P inhibits HK1-clustering, likely by binding to the active site of the C-terminal half,^9^ stabilizing a closed conformation.^28^ Since we found that monomeric HK1 favors HK1-clustering, we propose that dimer-monomer-transition is the second step in forming HK1-clusters. However, we do not yet know whether this transition occurs and if it is necessary for the HK1-clustering. Another possibility is that the previously determined dimer interface is different from the one involved in HK1-clustering.^25^ We propose that oligomerization of monomeric HK1 is the final step in HK1-clustering. We found that the N-terminal half of HK1 is essential for HK1-clustering. Although the N-terminal halves of HK1 and HK2 have similar structural folds, only the N-terminal half of HK1 can support a partially open conformation.^29^ This structural detail may explain why the N-terminal half of HK2 cannot substitute for that of HK1. Our findings suggest that leucine residues within the MBD of HK1 are crucial for the formation of HK1-clusters. Although it is striking that mutations in the MBD can prevent HK1-clustering, it has been shown that mutations in membrane anchors hinder protein oligomerization by preventing tight packaging or the ability to sense curvature.^30,31^ Collectively, our data show that HK1-clustering only occurs during severe cellular energy stress, defined by the depletion of ATP and G6P. Intense exercise, ischemia, and stroke have been reported to induce severe energy stress,^32–34^ suggesting that HK1-clustering likely plays a role in these conditions.

Our findings shed light on the potential functions of HK1-rings. We found that HK1-rings are positioned at the ER-mitochondria contact sites. These sites play crucial roles in various cellular processes, including calcium homeostasis, lipid transfer, and mitochondrial metabolism.^35^ However, we do not yet know whether HK1-rings are involved in these processes or whether HK1-rings influence the communication between mitochondria and the ER during energy stress. We found that mitochondrial constriction occurred at positions of HK1-rings. Thus, HK1-rings may play a role in defining the position of mitochondrial constriction during energy stress. Although actin is known to contribute to the constriction of mitochondria,^12,13^ we found that actin polymerization is not required to form HK1-rings. HK1 and actin share structural similarities, indicating possible common ancestry.^36^ Although it was assumed that HK1 had lost its ability to polymerize and generate mechanical force,^37^ our findings show that HK1 polymerizes into rings that constrict mitochondria. Our data also demonstrate that severe constriction of mitochondria by HK1-rings displaces cristae membrane. Constriction of mitochondria and decreasing cristae density are initial steps of mitochondrial fission.^12,38^ However, we did not observe mitochondrial fission at HK1-rings. Instead, we found that HK1-rings prevented mitochondrial fission when mitochondria were challenged with a chemical uncoupler. The inhibition of mitochondrial fission by HK1-rings was likely caused by the inability of Drp1 to localize at positions of HK1-rings. HK1-rings appeared to be dense structures capable of displacing outer mitochondrial membrane proteins. At a few sites of HK1-ring disassembly, we found that Drp1 clustered and triggered mitochondrial fission. Thus, we propose that HK1-rings prevent mitochondrial fission during energy stress but likely promote fission in some mitochondria after restoration of cellular energy. The importance of promoting fission after restoration of cellular energy remains unclear. However, inhibition of mitochondrial fission is known to reduce apoptosis and mitochondrial degradation.^39,40^ Therefore, we believe that inhibition of mitochondrial fission by HK1-rings can protect against excessive fission in ischemia/reperfusion injury and heart failure.^41^ We found that HK1-rings shift cellular metabolism towards glutaminolysis and increased TCA cycle activity. We think that these alterations could be linked to the constriction of mitochondria by HK1-rings, since changes in mitochondrial morphology have been shown to affect the sensitivity of enzymes to substrates.^42^ However, further research is needed to understand the mechanisms of how HK1-rings rewire cellular metabolism.

## Limitations of this study

It would be interesting to investigate HK1-rings in living animal models during severe energy stress such as ischemia or stroke. It is also important to understand the long-term effects and interaction partners of HK1-rings. Although we could not identify the crystal structure of HK1-polymers, we predict that studying full-length HK1 without G6P and ATP has a high chance of revealing the organization of HK1-clusters. Moreover, future studies with larger sample sizes and different biological samples are needed to further substantiate our NMR metabolomics findings. We do not yet know enough about the physiological and pathological relevance of our findings but they have a wide range of implications for how HK1-rings affect mitochondrial functions during energy stress.

## Methods

### Cell culture

HeLa, SH-SY5Y, MCF7, and MEF cells were cultured in Dulbecco’s modified Eagle’s medium (DMEM D5523, Sigma-Aldrich) supplemented with 10% FCS, 10 mM NaHCO_3_, 50 U/mL penicillin-streptomycin, 1.25 µg/mL amphotericin B and 25 mM HEPES; pH was adjusted to 7.45 with NaOH. INS-1 cells were cultured in GIBCO RPMI medium 1640 supplemented with 10% FCS, 10 mM HEPES, 2 mM L-glutamine, 1 mM sodium pyruvate, 0.05 mM beta-mercaptoethanol, 50 U/mL penicillin-streptomycin, 1.25 µg/mL amphotericin B. All cells were grown in a humified atmosphere of 5% CO_2_ at 37°C.

### Transfection

Cells were seeded in 6-well plates on 30 mm glass coverslips (Paul Marienfeld GmbH & Co. KG, Lauda-Königshofen, Germany) and transfected using PolyJet (SignaGen Laboratories). Per well, 3 µl of PolyJet reagent was mixed with 1 µg of plasmid DNA in 100 µl of DMEM devoid of serum and antibiotics. The transfection mixture was added to 1 ml of culture medium for 8 hours and was then replaced with 2 ml of culture medium. Imaging was performed 24 - 48 h after transfection.

### Imaging of subcellular protein dynamics

Before imaging, cells were put into a storage buffer, which was composed of 138 mM NaCl, 5 mM KCl, 2 mM CaCl_2_, 1 mM MgCl_2_, 10 mM HEPES, 2.6 mM NaHCO_3_, 0.44 mM KH_2_PO_4_, 0.34 mM Na_2_HPO_4_, 10 mM D-glucose, 2 mM L-glutamine, 1X MEM amino, 1X MEM vitamins, 1% penicillin-streptomycin and 1% Amphotericin B; pH was adjusted to 7.45 with NaOH.

High-resolution imaging was performed with an array confocal laser scanning microscope (Axiovert 200 M, Zeiss) equipped with a 100×/1.45 NA oil immersion objective (Plan-Fluor, Zeiss) and a Nipkow-based confocal scanner unit (CSU-X1, Yokogawa Electric Corporation). Laser light of diode lasers (Visitron Systems, Pucheim, Germany) served as the excitation light source: BFP, GFP, and RFP fusion constructs were excited with 405, 488, and 561 nm lasers, respectively. Emission light was captured with a CoolSNAP HQ2 CCD Camera (Photometrics Tucson, Arizona, USA) using the emission filters ET460/50m, ET525/36m, and ET630/75m (Chroma Technology Corporation) for BFP, GFP, and RFP fusion constructs, respectively.

Super-resolution imaging was performed with a structured illumination microscope (Nikon) equipped with a 100×/1.49 NA oil immersion objective (CFI Aopchromat TIRF, Nikon), standard filter sets, and two iXon EMCCD cameras (Andor). GFP and RFP fusion constructs were excited with 488 and 561 nm lasers, respectively.

Image analysis was performed with Fiji software. Z-stack images with a step size of 200 nm were deconvoluted and background-subtracted using a rolling ball radius of 50 to 300 pixels. For colocalization analysis, ROIs were drawn around cells using the polygon tool, and the Pearson coefficient was measured using the ImageJ plugin coloc2. The width of mitochondria was measured with the full width at half maximum. The TrackMate plugin was used to quantify HK1-clusters, HK1-rings, and Drp1-clusters. To assess mitochondrial morphology, 2D images of a mitochondrial matrix marker were thresholded using the Otsu method, and the ratio of the longest and shortest axes of a mitochondrial matrix marker was calculated.

### Single-cell ATP and glucose imaging

ATP and glucose FRET-imaging was performed with an inverted microscope (IX73 system, Vienna, Austria) equipped with a 40×/1.4 NA oil immersion objective (UPLXAPO40XO, Olympus) and an optical beam splitter (Photometrics DV2, Photometrics, Tucson, AZ, USA). HeLa cells expressing the mitochondria-targeted mtAT1.03 or cytosolic FLII12Pglu-700uDelta6 CFP-YFP-based biosensors were imaged. To excite the CFP of the FRET pair, we used an LED-based light source (LEDHub, Omicron) and a 427/10 Brightline HC excitation filter (Semrock). Emission light was captured with a Retiga R1 CCD camera (Teledyne QImaging) using a CFP/YFP/mCherry triple LED HC emission filter (Semrock).

### Perfusion of cells during live cell imaging

Transfected cells on 30 mm coverslips were put in a PC30 perfusion chamber (NGFI, Graz, Austria) and perfused at a flow rate of approximately 1 ml per minute with a gravity-based perfusion system (PS9, NGFI). Glucose buffer was composed of 135 mM NaCl, 5 mM KCl, 2 mM CaCl_2_, 1 mM MgCl_2_, 10 mM HEPES, and 10 mM D-glucose; pH was adjusted to 7.45 with NaOH. The glucose-free buffer contained 10 mM D-mannitol instead of glucose.

### Cell permeabilization

Cells were perfused with 10 µM digitonin for 4 min in a buffer mimicking the intracellular ion composition. The buffer was composed of 10 mM NaCl, 110 mM KCl, 1 mM MgCl_2_, 0.1 mM EGTA, 10 mM HEPES, and 10 mM D-glucose; pH was adjusted to 7.4 with NaOH. The glucose-free intracellular buffer contained 10 mM D-mannitol instead of glucose. For dose-response experiments, ascending levels of ATP or G6P were added to the glucose-free intracellular buffer, and the percentage of cells with HK1-clusters was assessed. For phosphate addition experiments, 100 µM G6P was added to phosphate-buffered saline (PBS).

### Chemicals

To stain the inner mitochondrial membrane, cells were incubated with 200 nM MitoTracker Red CMXRos (Thermofisher) in loading buffer for 10 minutes at 37°C and washed afterward with loading buffer. To inhibit actin polymerization, cells were perfused with 10 µM cytochalasin D (Sigma) in glucose buffer. To induce mitochondrial fission, cells were perfused with 2 µM FCCP (Sigma) in glucose-free buffer.

### Immunofluorescence

MCF7 cells were seeded in 6-well plates on 30 mm coverslips to reach a confluency of ∼80%. Cells were then taken from the incubator and put in loading buffer containing 1 µM MitoTracker Red CMXRos for 10 min at 37°C. To induce HK1-ring formation, cells were perfused with glucose-free buffer for 30 minutes. Control cells were perfused with glucose buffer. All cells were washed twice with PBS and fixed with 4% paraformaldehyde at room temperature for 15 min. Then cells were washed 3x with PBS and blocked with 5% BSA in PBS for one hour while mildly shaking. Then the blocking buffer was replaced with 1:1000 diluted HK1 rabbit monoclonal antibody (cell signaling, #2024) in PBS with 5% BSA. Cells were incubated with the primary antibody overnight at 4°C, mildly shaking. On the next day, primary antibody solution was removed, cells were washed 3x with PBS, and secondary antibody solution containing Alexa Fluor 488 goat anti-rabbit (Thermo Fisher Scientific) at a dilution of 1:2.500 was added. Cells were incubated for 2 hours in the dark at room temperature, mildly shaking. Finally, cells were washed 3x for 5 minutes with PBS, Vectashield antifade mounting medium (Vectorlabs), and the coverslips were sealed with nail polish.

### Plasmids

| Name | Description | Reference |
| --- | --- | --- |
| HK1-GFP | HK1 marker | <sup>19</sup> Addgene #21917 |
| HK1-mRuby3 | HK1 marker | <sup>43</sup> |
| HK1-ATPmut-GFP | ATP binding mutant G862A<sup>11</sup> | This study |
| Monomeric-HK1-GFP | E280A, R283A, G284Y mutations prevent dimerization of HK1<sup>25</sup> | This study |
| HKDC1-GFP | Hexokinase domain containing 1 | This study |
| HK2-GFP | HK2 marker | <sup>19</sup> Addgene #21920 |
| HK1N-HK2C-GFP | The N-terminal half of HK1 fused to C-terminal half of HK2 | This study |
| HK2N-HK1C-GFP | N-terminal half of HK2 fused to C-terminal half of HK1 | This study |
| Truncated-HK1-GFP | Lacks mitochondrial binding domain (MBD) | <sup>19</sup> Addgene #21918 |
| HK1MBD-GFP | Mitochondrial binding domain of HK1 | This study |
| HK2MBD-HK1-GFP | Mitochondrial binding domain of HK1 was replaced by MBD of HK2 | This study |
| HK1-L6A-GFP | Mutation in the MBD | This study |
| HK1-L7A-GFP | Mutation in the MBD | This study |
| HK1-F11A-GFP | Mutation in the MBD | This study |
| HK1-L14A-GFP | Mutation in the MBD | This study |
| mitoDsRed | Mitochondrial matrix marker | Addgene #44386 |
| mito-sfGFP | Mitochondrial matrix marker | <sup>44</sup> |
| mitoBFP | Mitochondrial matrix marker | Addgene #49151 |
| ComplexIV8-mRuby2 | Cristae membrane marker | <sup>45</sup> |
| BFP-KDEL | ER lumen marker | Addgene #49150 |
| mCherry-Actin | Actin marker | Addgene #54957 |
| mCherry-Drp1 | Drp1 marker | Addgene #49152 |
| BFP-Tom20 | Tom20 marker | Addgene #55328 |
| mtAT1.03 | Mitochondrial matrix targeted FRET-based ATP sensor | <sup>7</sup> |
| FLII12Pglu-700uDelta6 | Cytosolic FRET-based glucose sensor | <sup>8</sup> Addgene #17866 |

## Acknowledgments

We appreciate the technical assistance from René Rost, Anna Schreilechner, Mercedes Maier, and Sinem Usluer. The research was supported by the Molecular Medicine PhD program of the Medical University of Graz and the FWF (Austrian Science Fund: I3716-B27 to R.M.). We thank Emil Spreitzer for his insightful analysis. We appreciate all discussions about this work, especially with Marc Germain and Marijn Ford. We thank everyone who sent us plasmids or deposited plasmids used in this study at www.addgene.org. We are grateful to the reviewers and editors for their precious time.

## Author Contributions

J.P. and R.M. conceived the study. J.P. and B.G. performed experiments and carried out image analysis. H.H. carried out NMR metabolomics. All authors contributed to the interpretation of results and experimental design. J.P. and R.M. wrote the first draft of the manuscript, and all authors contributed to the final version.

