## Supplementary Figures for "Hexokinase 1 forms rings that constrict mitochondria during energy stress"

### Figure S1. Hexokinase 1 clusters into ring-like structures during energy stress

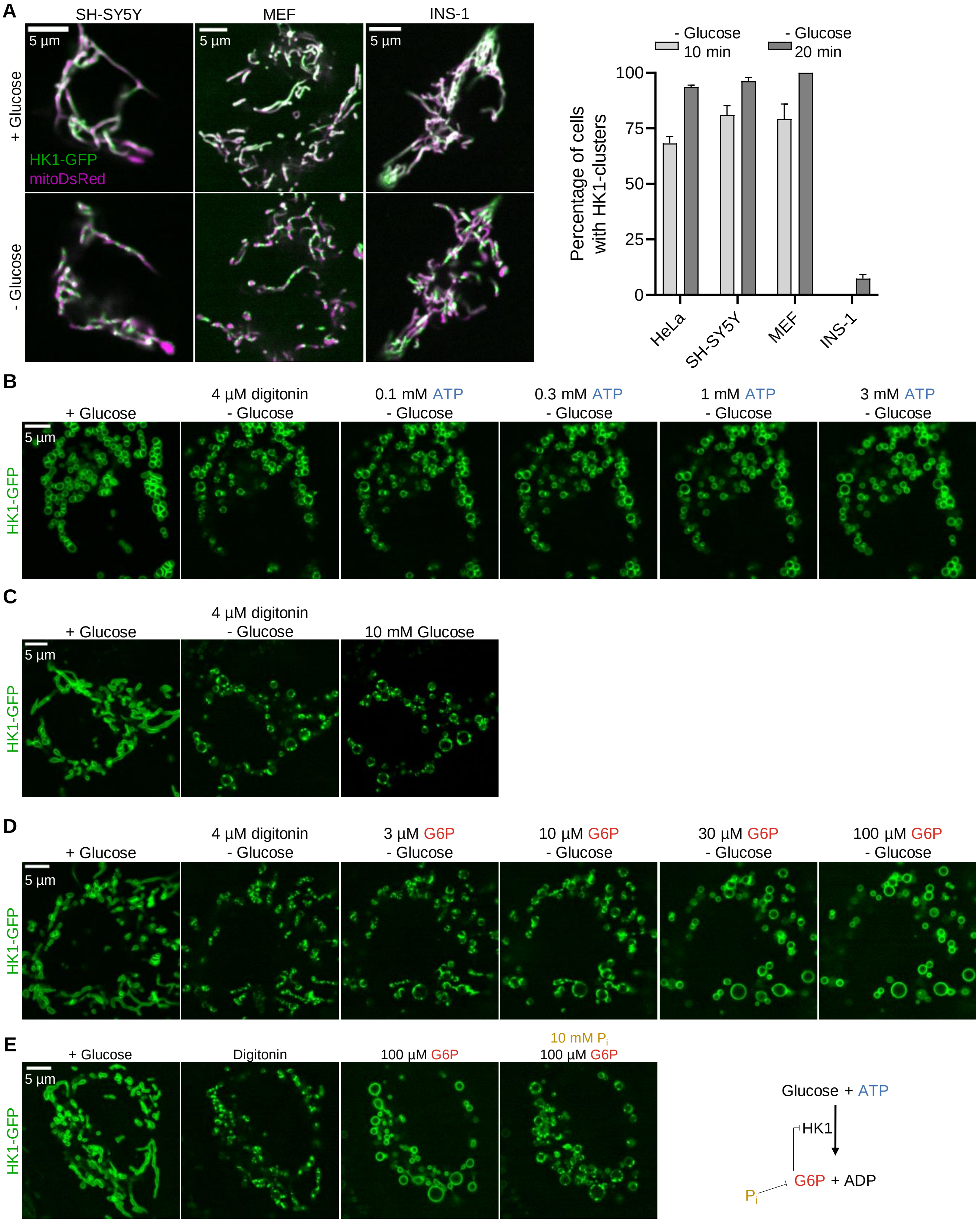

### Figure S1. Hexokinase 1 clusters into ring-like structures during energy stress

**F**

Percentage of cells with clusters

| | 10 mM Glc | 1 mM Glc | 0.33 mM Glc | 0 mM Glc |
|---|---|---|---|---|
| HK1 | 0% | 0% | 22% | 93% |
| HK1-ATPmut | 2% | 15% | 79% | 98% |

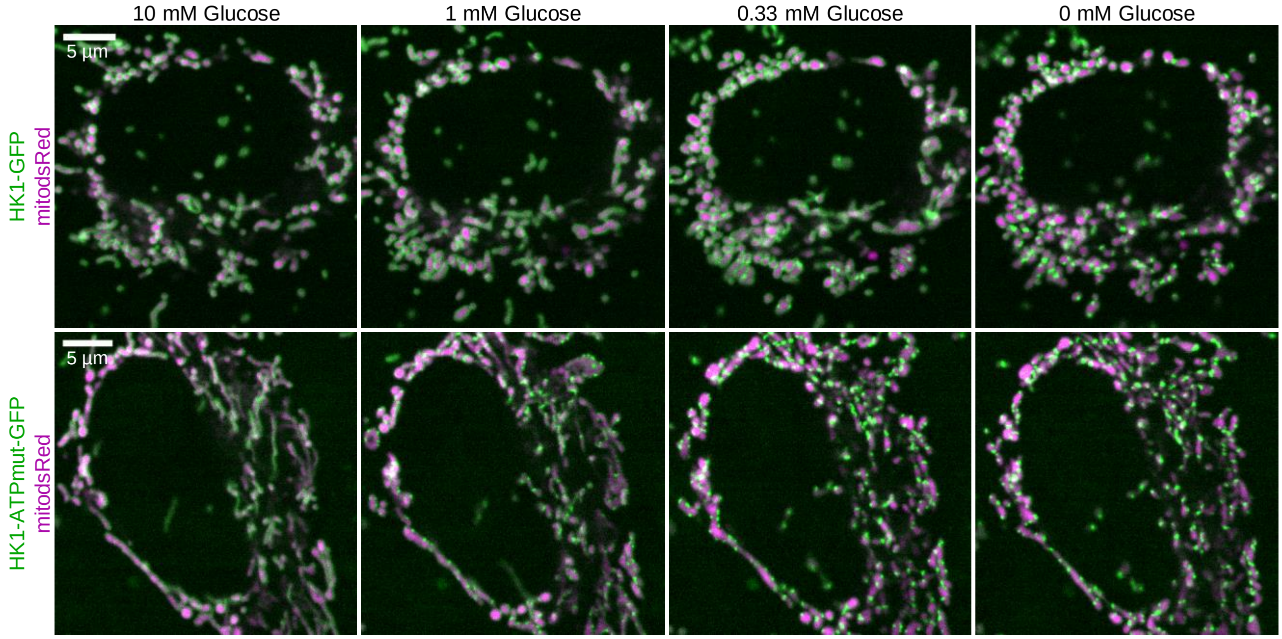

### Figure S2. HK1-rings constrict mitochondria at-ER contact sites

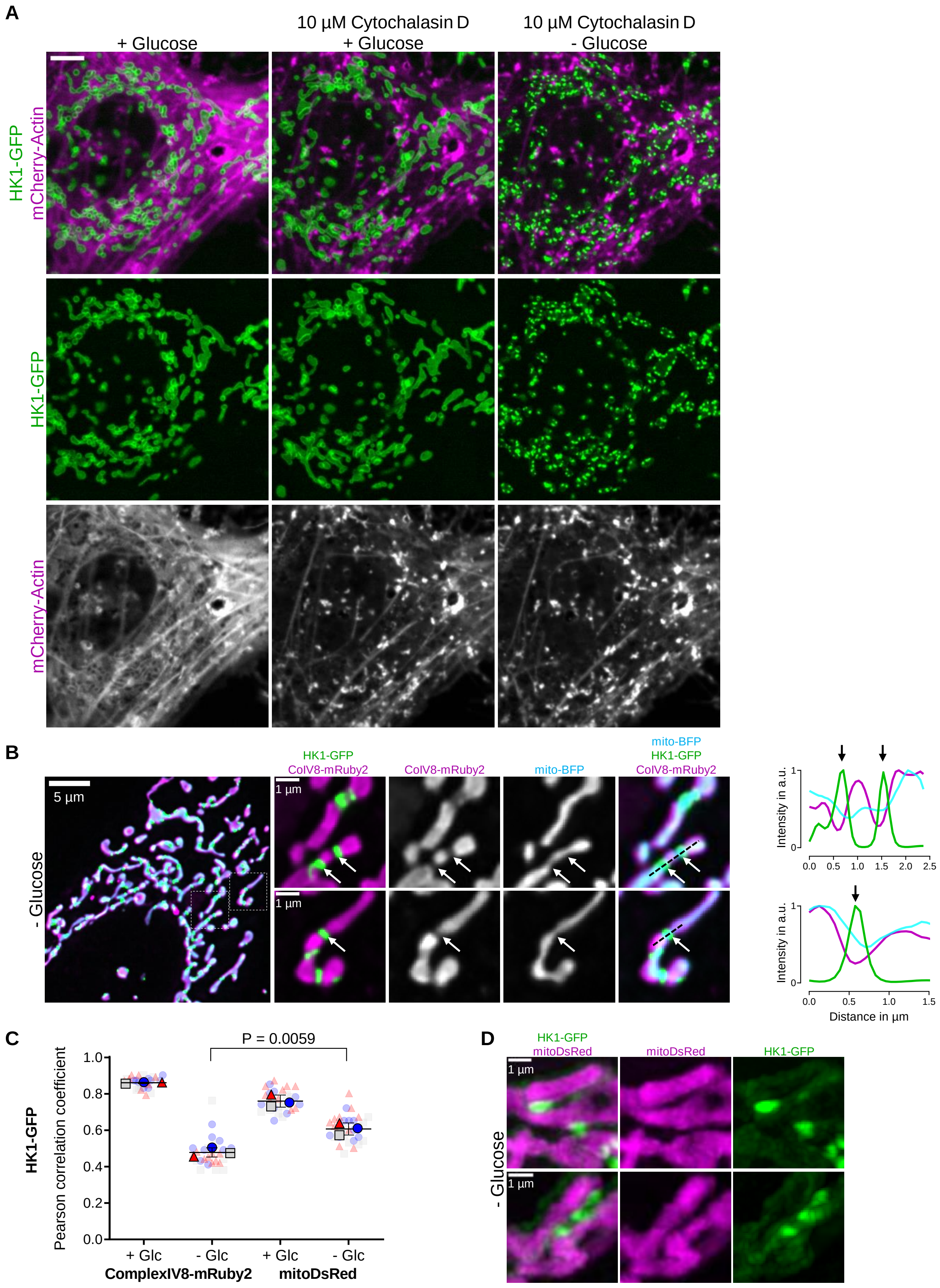

### Figure S3. HK1-rings displace Drp1 and prevent mitochondrial fission during energy stress

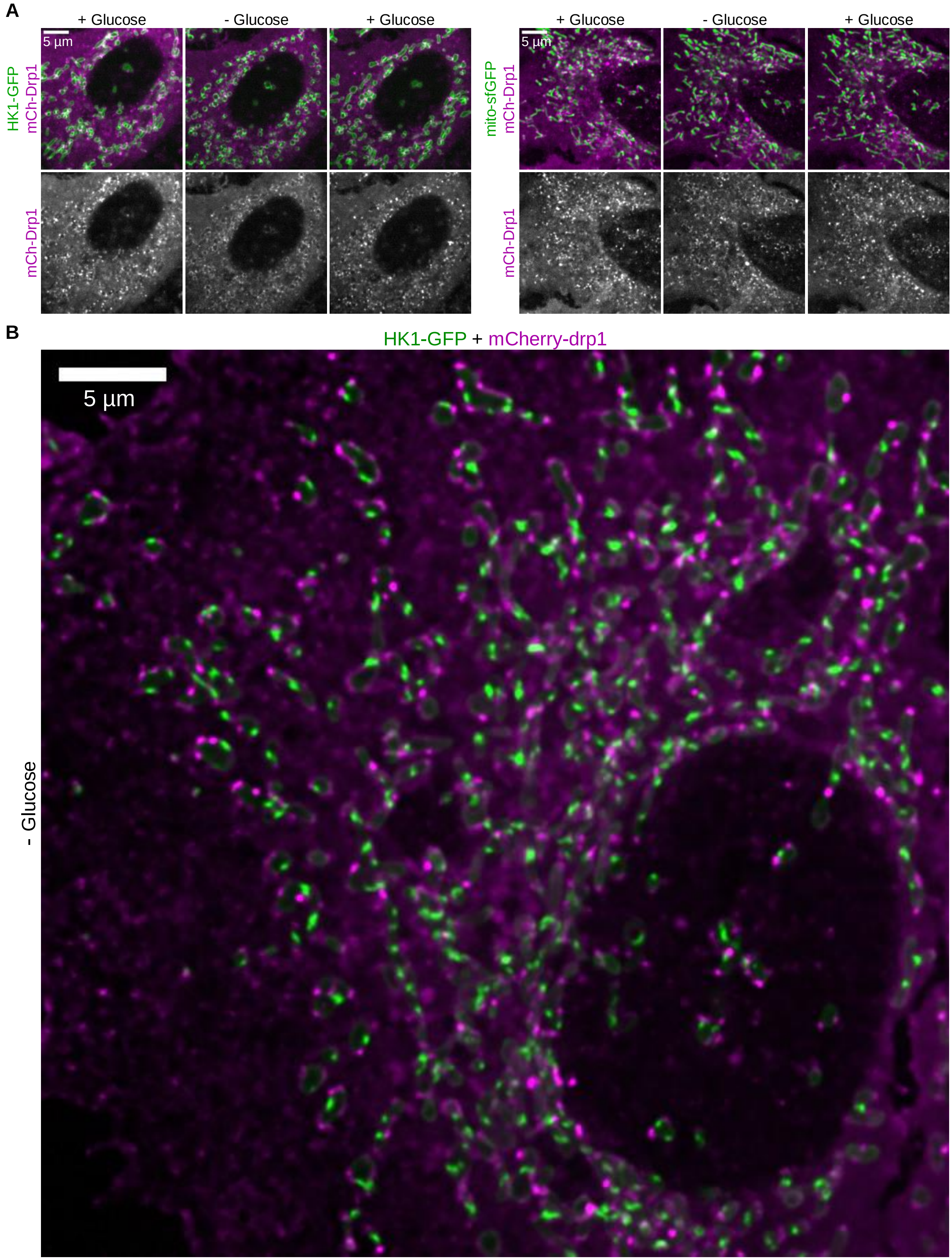

### Figure S4. Structural features dictate the formation of HK1-clusters

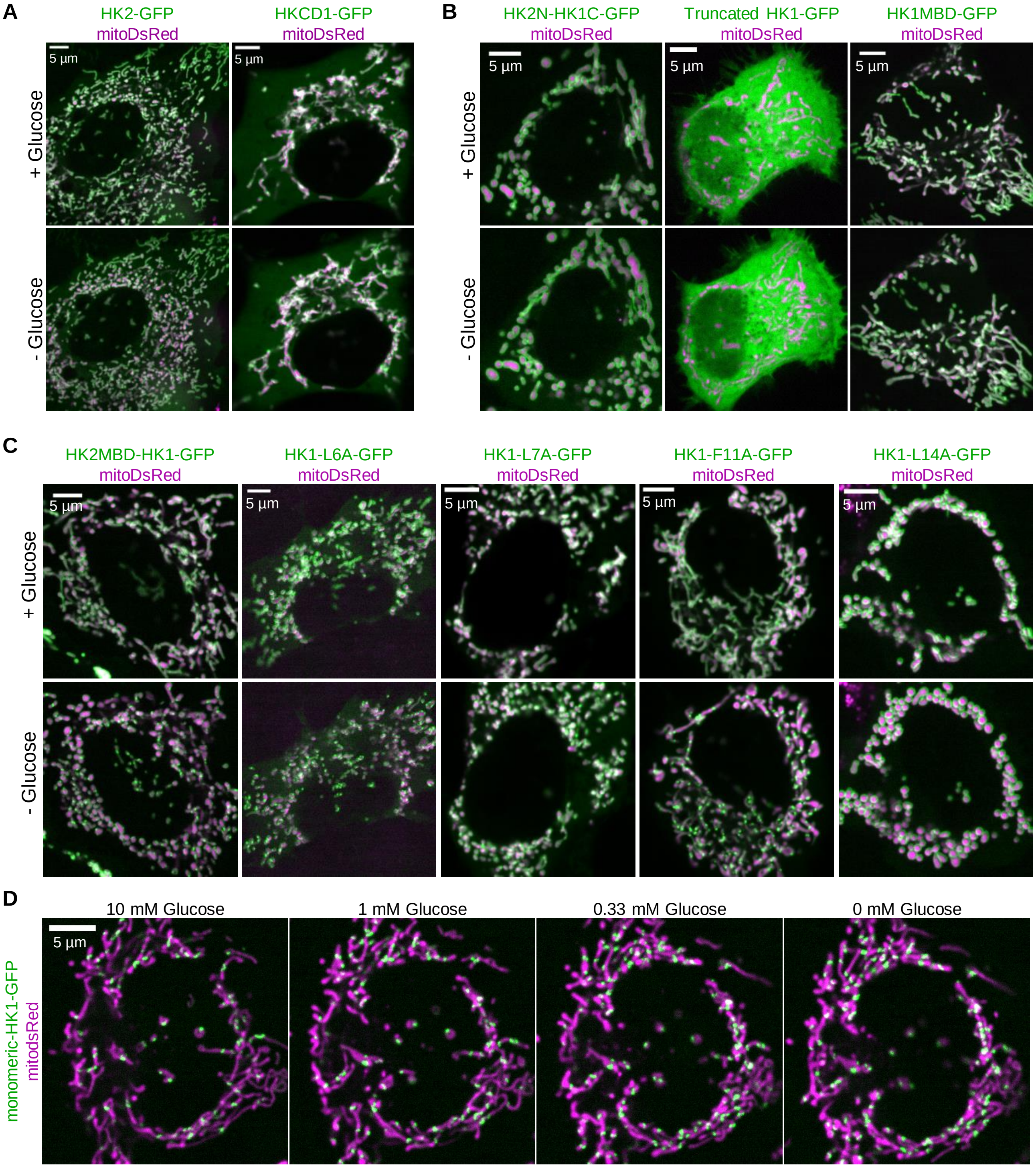

**Figure S5. HK1-rings keep mitochondria connected and rewire cellular metabolism**

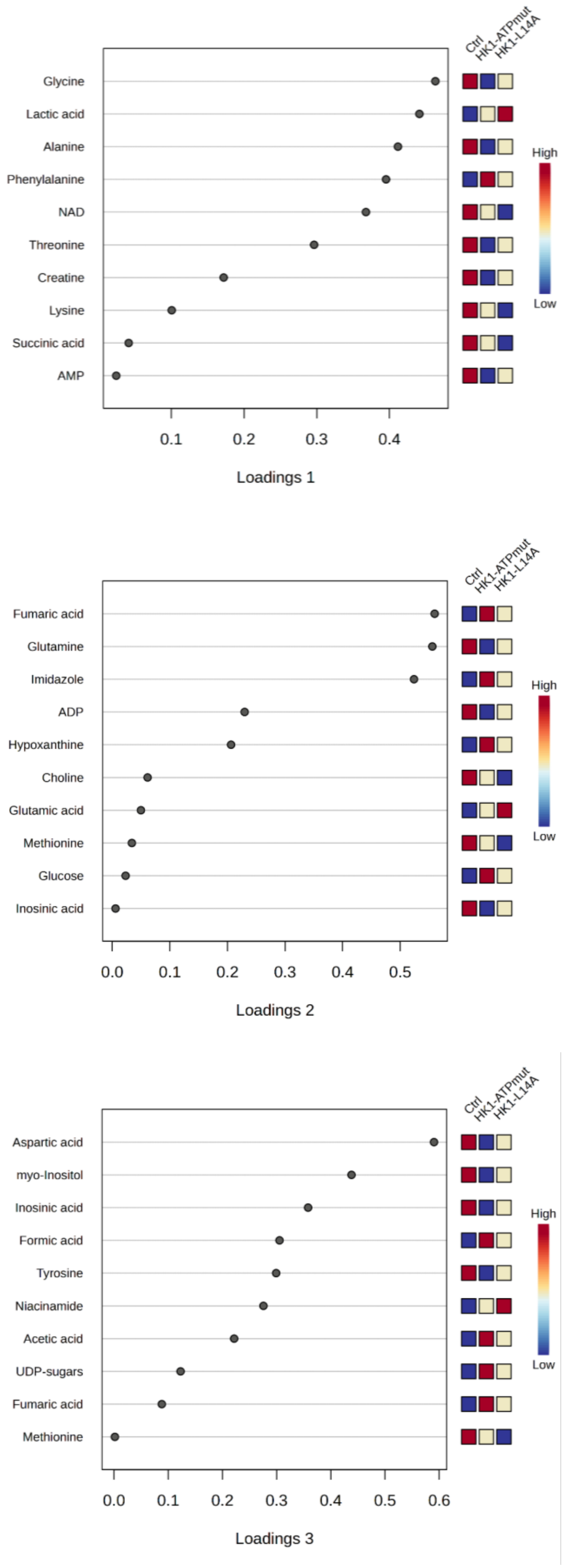
