## Supplementary Information for "Hexokinase 1 forms rings that constrict mitochondria during energy stress"

**Supplementary figure legends**

**Figure S1. Hexokinase 1 clusters into ring-like structures during energy stress**

(A) HK1-clustering occurs in various cell types. The left panel shows confocal images of SH-SY5Y, MEF, and INS-1 cells expressing HK1-GFP and mitoDsRed with 10 mM glucose (top) and without glucose (bottom). The right bar graph shows the percentages of cells with HK1-clusters (mean ± SEM) as glucose is depleted for 10 min (light gray) or 20 min (dark gray). Number of cells: HeLa (n = 110), SH-SY5Y (n = 53), MEF (n = 24), INS-1 (n = 54).

(B-D) ATP and glucose-6-phosphate (G6P) disassemble HK1-clusters in a concentration-dependent manner. Time-lapse images of HK1-GFP-expressing HeLa cells permeabilized with 4 µM digitonin before being treated with ATP (B), glucose (C), or G6P (D). Time-lapse images were acquired at intervals of 7 min.

(E) Phosphate (P_i_) promotes HK1-clustering despite the presence of G6P. Time-lapse images of HK1-GFP-expressing HeLa cells permeabilized with 4 µM digitonin before being treated with G6P and P_i_. Time-lapse images were acquired at intervals of 7 min. The circuit on the right shows the regulation of the HK1 reaction.

(F) The ATP-binding mutant (G862A) is more likely to form clusters than the wild-type enzyme. Time-lapse images of HeLa cells expressing mitoDsRed and HK1-GFP or HK1-ATPmut-GFP were acquired at intervals of 7 min as glucose was gradually depleted (bottom). The percentages of cells with clusters of HK1 (n = 41) and HK1-ATPmut (n = 47) were calculated (top).

**Figure S2. HK1-rings constrict mitochondria at ER contact sites**

(A) The formation of HK1-rings does not require actin polymerization. Time-lapse images of a HeLa cell expressing HK1-GFP and mCherry-Actin were acquired at intervals of 10 min under three different conditions: With 10 mM glucose (left), after treatment with 10 µM cytochalasin D (middle), and after glucose depletion in the presence of 10 µM cytochalasin D (right).

(B) Mitochondrial matrix is present in cristae-free regions at positions of HK1-rings. Confocal images of a HeLa cell expressing HK1-GFP, ComplexIV8-mRuby2 (cristae membrane), and mitoBFP (matrix) after 15 min of glucose depletion. Images are maximum intensity projections of z-stacks (36 sections, spaced 0.2 µm apart). The left image shows an overview of the cell, and the dashed squares are magnified on the right side. Dashed lines represent line scan graphs on the right and show the relative fluorescence intensity of HK1 (green), cristae membrane (magenta), and mitochondrial matrix (blue) along the length of the line. Arrows point to positions of HK1-rings.

(C) HK1-rings colocalize less with cristae membrane than with mitochondrial matrix. The Pearson correlation coefficient is shown between HK1-GFP and ComplexIV8-mRuby2 (cristae membrane), or mitoDsRed (matrix) with 10 mM glucose or after 15 min of glucose depletion. The beeswarm SuperPlot represents each cell with a color-coded dot according to the experimental day. Two-tailed unpaired t-test was used for statistical analysis (n = 3). Data are presented as mean ± SD.

(D) HK1-clusters of neighboring mitochondria were frequently observed in contact. Structured illumination microscopy images of HeLa cells expressing HK1-GFP and mitoDsRed, showing that HK1-clusters from neighboring mitochondria are in lattice contacts. Images are maximum intensity projections of z-stacks (33 sections, spaced 0.1 µm apart).

**Figure S3. HK1-rings displace Drp1 and prevent mitochondrial fission during energy stress**

(A) Overexpression of HK1 significantly reduced the number of Drp1-clusters during energy stress. Confocal images of HeLa cells expressing mCherry-Drp1 with HK1-GFP (left panel) or mito-sfGFP (right panel) as 10 mM glucose (left) was removed for 20 min (middle) and readded for 4 min (right).

(B) The majority of HK1-rings and Drp1-clusters did not colocalize. Representative confocal image of a HeLa cell expressing HK1-GFP and mCherry-Drp1 after 15 min of glucose depletion. The image is a maximum intensity projection of z-stacks (41 sections, spaced 0.2 µm apart).

**Figure S4. Structural features dictate the formation of HK1-clusters**

(A) HK2 does not form clusters, and HKCD1 has a moderate ability to form clusters when glucose is depleted. Confocal images of HeLa cells expressing mitoDsRed and HK2-GFP (left) or HKCD1-GFP (right) with 10 mM glucose (top) and after 10 min of glucose depletion (bottom).

(B) HK2N-HK1C, truncated HK1, and HK1MBD do not form clusters when glucose is depleted. Confocal images of HeLa cells expressing mitoDsRed and HK2N-HK1C-GFP (left), truncated HK1-GFP (middle), or HK1MBD-GFP (right) with 10 mM glucose (top) and after 10 min of glucose depletion (bottom). MBD, mitochondrial binding domain; N, N-terminal half; C-terminal half.

(C) Leucine residues within the MBD are crucial for the formation of HK1-clusters. Confocal images of HeLa cells expressing mitoDsRed and HK2MBD-HK1-GFP, HK1-L6A-GFP, HK1-L7A-GFP, HK1-F11A-GFP, and HK1-L14A-GFP with 10 mM glucose (top) and after 10 min of glucose depletion (bottom).

(D) Monomeric HK1 (E280A, R283A, and G284Y) forms clusters even in the presence of glucose. Time-lapse images of a HeLa cell expressing or monomeric HK1-GFP and mitoDsRed were acquired at intervals of 7 min as glucose was gradually depleted.

**Figure S5. HK1-rings keep mitochondria connected and rewire cellular metabolism**

HK1-rings rewire mitochondrial metabolism. Loading plots indicate the most distinctive metabolites. The tables with the listed metabolites show changes in concentration. Heatmaps are shown on the right side.

**Supplemental video legends**

**Video S1A. Formation of HK1-clusters**

Time-lapse images of a HeLa cell expressing HK1-GFP and mitoDsRed were acquired at intervals of 90 s as glucose was depleted.

**Video S1B. 3D animation of an HK1-cluster**

Three-dimensional animation of an HK1-cluster in a HeLa cell expressing HK1-GFP and mitoDsRed. Z-stack images were acquired with structured illumination microscopy (33 sections, spaced 0.1 µm apart).

**Video S1C. 3D animation of an HK1-ring**

Three-dimensional animation of an HK1-ring in a HeLa cell expressing HK1-GFP and mitoDsRed. Z-stack images were acquired with structured illumination microscopy (28 sections, spaced 0.1 µm apart).

**Video S1D. Disassembly of HK1-clusters**

Time-lapse images of a glucose-depleted HeLa cell expressing HK1-GFP and mitoDsRed were acquired at intervals of 5 s as 10 mM glucose was readded.

**Video S2A. 3D animation of HK1-ring interaction with ER**

Three-dimensional animation of HK1-ring interaction with ER in a HeLa cell expressing HK1-GFP, mitoDsRed, and BFP-KDEL (ER). Z-stack images were acquired with confocal microscopy (37 sections, spaced 0.2 µm apart).

**Video S2B. 3D animation of HK1-cluster interaction with ER**

Three-dimensional animation of HK1-cluster interaction with ER in a HeLa cell expressing HK1-GFP, mitoDsRed, and BFP-KDEL (ER). Z-stack images were acquired with confocal microscopy (37 sections, spaced 0.2 µm apart).

**Video S3A. HK1-rings prevent mitochondrial fission**

Time-lapse images of HeLa cells stained with mitoTracker (magenta) were acquired at intervals of 1 min during glucose depletion. The cell on the left shows no HK1-GFP expression while HK1-rings are visible in the cell on the right.
